# Semi-supervised integration of single-cell transcriptomics data

**DOI:** 10.1101/2023.07.07.548105

**Authors:** Massimo Andreatta, Léonard Hérault, Paul Gueguen, David Gfeller, Ariel J Berenstein, Santiago J Carmona

## Abstract

Single-cell sequencing technologies offer unprecedented opportunities to characterize the complexity of biological samples with high resolution. At the same time, variations in sample processing and experimental protocols introduce technical variability – or “batch effects” – in the molecular readouts, hindering comparative analyses across samples and individuals. Although batch effect correction methods are routinely applied in single-cell omics analyses, data integration often leads to overcorrection, resulting in the loss of true biological variability. In this study, we present STACAS v2, a semi-supervised scRNA-seq data integration method that leverages prior knowledge in the form of cell type annotations to preserve biological variance. Through an open and reproducible benchmarking pipeline, we show that semi-supervised STACAS outperforms popular unsupervised methods such as Harmony, FastMNN, Seurat v4, scVI, and Scanorama, as well as supervised methods such as scANVI and scGen. Notably, STACAS is robust to incomplete and imprecise cell type annotations, which are commonly encountered in real-life integration tasks. Highlighting its scalability, we successfully applied semi-supervised STACAS to construct a high-resolution map of tumor-infiltrating CD8 T cells encompassing over 500,000 cells from 265 individuals. Based on our findings, we argue that the incorporation of prior cell type information should be a common practice in single-cell data integration, and we provide a flexible framework for semi-supervised batch effect correction. STACAS seamlessly integrates with Seurat pipelines and can be run with one command: Run.STACAS(seurat.list, cell.labels).

## Introduction

Single-cell omics technologies enable characterizing the cellular complexity of biological samples with very high resolution. While individual samples can provide readouts for thousands of individual cells, addressing biological questions typically requires the comparative analysis of multiple samples, tissues, individuals and experimental conditions. By means of data integration or harmonization, cells from different sources can be placed in the same embedding or latent space, facilitating the measurement of distances between them, the collective annotation of cell populations, and additional downstream joint analyses. However, variations in sample collection, processing, and experimental protocols introduce unwanted variation in the molecular readouts that interferes with the identification of true biological differences between samples. This technical variation is sometimes referred to as “batch effects” since it is typically observed between groups of samples that were handled in different batches^1,2^.

Several methods for single-cell RNA-seq data integration have been proposed, based on different approaches such as mutual nearest neighbors and linear embeddings, deep learning, and graph structures, each with strengths and limitations^3–6^. Integration methods aim at removing batch effects while preserving relevant biological variation. Two main aspects are considered to determine the quality of single-cell data integration: *i)* batch mixing and *ii)* preservation of biological variance. Batch mixing measures whether similar cells from different batches are well mixed after integration. Frequently used metrics of batch mixing are entropy, kBET, and integration LISI (iLISI)^7–9^. Preservation of biological variance can be quantified by how close to each other cells of the same type are, and how separated from each other cells of different types are in the joint integrated embeddings. Commonly-used metrics include average silhouette width (ASW), average Rand index (ARI), and cluster LISI (cLISI)^9–11^. For a review on integration metrics see Luecken et al.^12^.

For certain tasks, such as integration of technical replicates or very similar samples, most integration methods perform generally well both in terms of batch mixing and preservation of biological variance^2,12^. However, more common scenarios include the integration of datasets from biologically heterogeneous samples, e.g. from different donors, timepoints or tissues. These do not only display technical batch effects, but also large variability in terms of cell type composition. Differences in cell type abundance are a major component of biological variance across samples^13,14^, and such imbalance in cell type composition makes datasets particularly susceptible to loss of biological variance during integration^15,16^.

The choice of a specific integration method and parameters configuration should take into account the tradeoff between preserving relevant biological variation and increasing batch mixing^12^. Since most integration tasks involve samples with some degree of cell type imbalance, choosing integration strategies that favor preservation of biological variance over batch mixing might be preferable^15^. An appealing approach to preserve biological variance is to make use of prior cell type information to guide dataset integration. Indeed, in recent benchmarks of single-cell data integration tools, methods that take cell type labels as input showed the highest performance in terms of preservation of biological variance^12,17^.

In this study, we describe STACAS v2, a semi-supervised scRNA-seq data integration method that leverages prior knowledge in the form of cell type annotations to preserve biological variance during integration. Using an open and reproducible benchmarking pipeline we show that semi-supervised STACAS compares favorably to popular unsupervised methods such as Harmony, FastMNN, Seurat v4, scVI, and Scanorama, as well as to supervised methods scANVI and scGen, while being robust to missing and imperfect cell type information. We argue that prior cell type information should be routinely incorporated in integration tasks and we propose a general strategy for its implementation.

## Results

### Semi-supervised STACAS uses prior cell type information to guide data integration

STACAS is a batch correction method for the integration of heterogeneous scRNA-seq datasets. The result of STACAS integration is a batch-corrected combined gene expression matrix that can be used for downstream multi-sample analyses, such as clustering and visualization. The method is based on the concept of mutual nearest neighbors to identify biologically equivalent cells in pairs of datasets (referred to as “anchors”)^3^, which are used to estimate batch effects. STACAS builds upon the Seurat integration method^5^ and appliesreciprocal principal component analysis (rPCA) to find anchors, where each dataset in a pair is projected into the principal components (PC) space of the other. STACAS uses the rPCA distance between the two cells of an anchor to weigh the biological relevance of the anchor and ultimately its contribution to batch correction vectors (see Methods). In this way, anchor cells that are close to each other in the rPCA space will contribute more strongly to batch correction than distant anchor cells, which are transcriptionally more dissimilar and thus less likely to be biologically equivalent cells.

In anchor-based methods, obtaining an accurate set of anchors is critical for integration performance. STACAS v2 introduces the ability to use prior information, in terms of cell type labels, to refine the anchor set. We refer to this mode of dataset integration as “semi-supervised”. Cell type labels may be obtained from automated classifiers, manual annotation, multi-modal information or any other source. When provided, cell labels are used by STACAS to remove “inconsistent” anchors, composed of cells with different labels (**Fig. 1**). Note that missing labels are not penalized in this step, generalizing the method to partially annotated datasets. Finally, consistent integration anchors are used to calculate batch effect correction vectors between pairs of datasets, and their weighted anchor scores are used to construct an integration guide tree that will determine the order in which the datasets will be integrated (see Methods).

**Figure 1:**
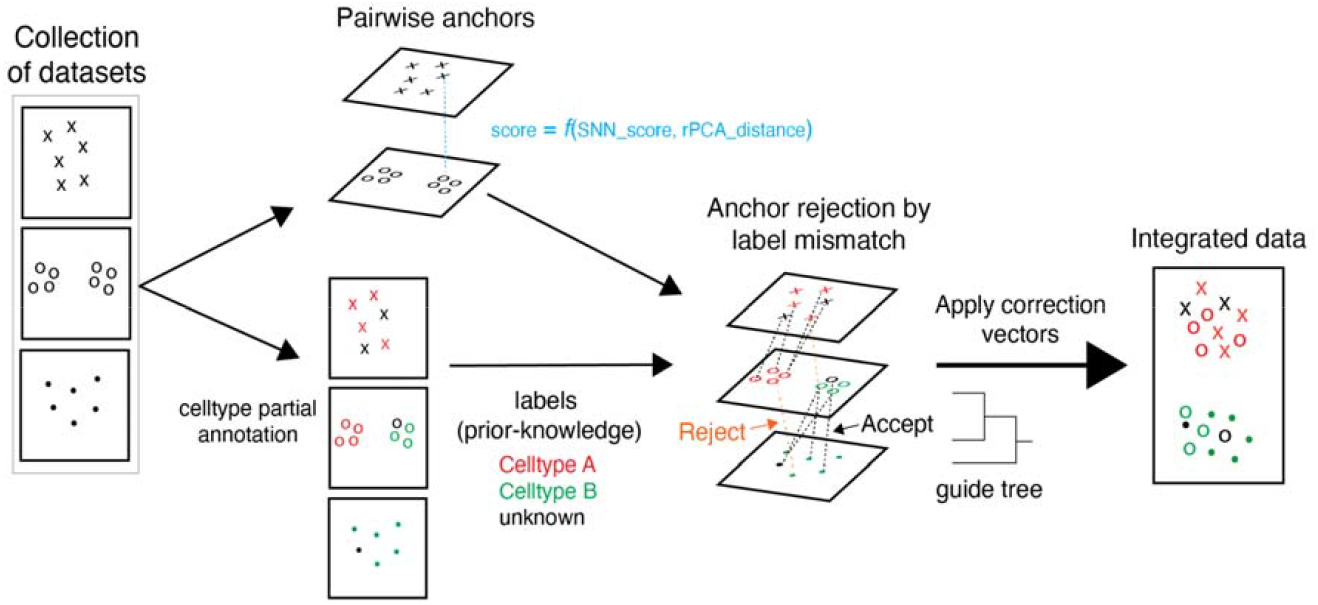
Schematic of (semi-supervised) STACAS integration method. The algorithm identifies integration anchors between all pairs of datasets from a shared nearest neighbors (SNN) graph. These are expected to be cells of the same type across batches and are used to calculate batch effects. Integration anchors are weighted by a score that combines a SNN anchor consistency score (based on the overlap of shared neighbors) and a score based on rPCA distance (how similar are cells of one dataset to the corresponding anchor cells in a second dataset projected into the PCA space of the latter). If cell type labels are available, they can be provided as input to the algorithm. When cell type labels between two cells of an anchor are inconsistent, the anchor is rejected with a predefined probability and in that case will not contribute to batch effect correction. Finally, the sum of retained, weighted integration anchor scores are used to calculate global similarities between datasets and to derive a guide tree that will determine the order in which the datasets are to be integrated.

### A cell type-aware implementation of the LISI metrics to quantify batch mixing

A frequently used metric to assess quality of single-cell data integration is the Local Inverse Simpson’s Index (LISI). LISI measures mixing by estimating the effective number of classes in local neighborhoods of cells^9^ and is relatively fast to compute. When applied to measure the effective number of datasets or batches in a neighborhood, this metric is referred to as ‘iLISI’ (integration LISI); when applied to measure the effective number of cell types in a neighborhood, LISI has been named ‘cLISI’ (“cluster” or “cell type” LISI). Another widely-used performance metric to assess cell type clustering is the average silhouette width (ASW), which quantifies distances of cells of the same type compared to the distances to cells of other types (cell type ASW) ^2,12^.

To evaluate the behavior of these metrics on datasets with different levels of cell type imbalance and batch effects, we generated synthetic scRNA-seq datasets (see Methods). This simulation setting consists of three biological samples: sample A comprises cell type 1 and cell type 2 in equal parts; sample B contains only cell type 2; and sample C contains only cell type 1. Each sample corresponds to a different experimental batch (batch 1 to 3) and displays batch effects in addition to cell type biological variation (**Fig. 2A**). We generated several scenarios with increasing batch effects: no batch effects (batch0), mild (batchMild), and strong batch effects (batchStrong). In the fourth example both cell type variation and batch effects are zero, representing the situation of an extreme overcorrection of batch effects. We employed these simulated datasets to evaluate different integration metrics.

**Figure 2:**
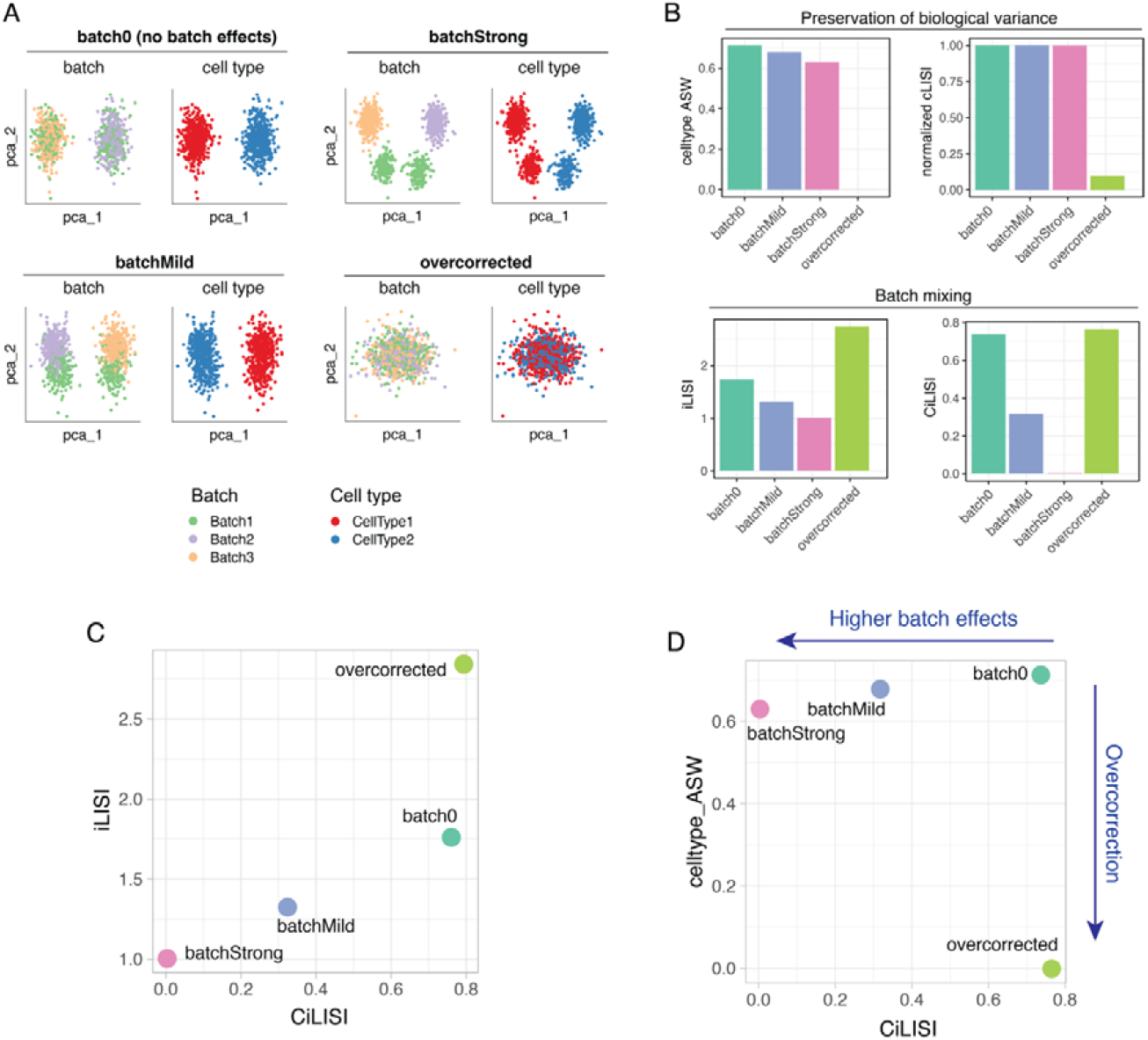
Integration metrics on synthetic single-cell datasets. **A)** Scatterplot of first and second principal components (pca_1 and pca_2) for four simulated scenarios where datasets have increasing levels of batch effects (“batch0” with no batch effects, “batchMild”, and “batchStrong”), and one case where there is no batch effect and no biological cell type signal (simulating the result of an extreme batch-effect “overcorrection”). Each scenario is composed of three samples/batches and two cell types. **B)** Metrics of batch mixing (average iLISI and average CiLISI) and of preservation of biological cell type variability (average cell type ASW and average normalized cLISI) for the five simulated scenarios. **C)** Scatterplot of average CiLISI (x-axis) versus average iLISI (y-axis) for the five simulated scenarios. **D)** Scatterplot of average CiLISI (x-axis) versus average cell type ASW (y-axis) for the five simulated scenarios; a batch-corrected and biological variance-preserving integration should aim at maximizing both metrics. CiLISI is defined here as the normalized batch LISI calculated on a per-cell type basis; it tends to 1 when cells of the same type are well mixed across batches, and becomes zero when cells from different batches do not mix.

In terms of preservation of biological variance, both normalized cell type ASW and normalized cLISI (see methods) correctly capture the poor cell type separation in the ‘overcorrected’ dataset, while remaining high in all other cases (**Fig. 2B**). We note that the normalized cLISI, because it only measures local neighborhoods, is unaffected by mild levels of batch effect. Instead, cell type ASW seems to be more sensitive in detecting cell type spread due to batch effects (**Fig. 2B**). In terms of batch mixing, iLISI decreases together with the mixing of cells from different batches, as expected. However, iLISI increases from ∼1.75 in the case with no batch effect (batch0) to ∼2.75 in the case with no batch effect and no cell type signal (overcorrected) (**Fig. 2B**). Hence, iLISI would favor a method that completely removes biological variance together with batch effects over a method that effectively removes batch effects while preserving biological variance. This is an undesirable behavior for an integration metric. To obviate this limitation, we propose to evaluate iLISI on a per-celltype basis, hereafter referred to as CiLISI (normalized to vary between 0 and 1, see Methods). Unlike iLISI, CiLISI measures batch mixing in a cell type-aware manner and scores similarly the cases with no batch effects, irrespective of the biological variance (**Fig. 2C**). We argue that CiLISI is preferable over iLISI because it does not penalize methods that preserve biological variance in datasets with cell type imbalance.

Given these insights, we suggest evaluating integration performance by assessing jointly *i)* batch mixing in terms of per-cell type normalized batch LISI (CiLISI) and *ii)* cell type clustering in terms of cell type ASW (or alternatively normalized cLISI). Well-performing methods should be able to mix cells of the same type in different batches (i.e. maximize CiLISI) while keeping apart cells of different types (i.e. maximizing cell type ASW and normalized cLISI) (**Fig. 2D**). We implemented these metrics in an R package, available at https://github.com/carmonalab/scIntegrationMetrics.

### Semi-supervised STACAS outperforms state-of-the-art methods

To assess the performance of STACAS v2 compared to state-of-the-art integration tools, we took advantage of the ‘scib’ pipeline published by Luecken et al.^12^, which allows comparing methods written in R and python in a reproducible environment. We modified the pipeline to include STACAS v2 and the CiLISI metric, and to evaluate all methods on latent spaces of the same size (see Methods for details). Importantly, we included the option to evaluate semi-supervised methods with incomplete cell type labels, as well as with partially shuffled labels. In our benchmark we shuffled 20% of the cell type labels and set 15% to ‘unknown’ as input to supervised methods, simulating a realistic setting where prior cell type knowledge is incomplete and imperfect. We assess the effect of overfitting in different integration methods as a function of the percentage of incomplete or noisy input labels in a later section.

The performance of 11 computational tools was evaluated on 4 different integration tasks, with increasing levels of cell type imbalance (**Fig. 3A-D**). We also evaluated the effect of re-scaling gene expression data prior to batch effect correction, and we report results for both unscaled and scaled data for each method (this is not applicable to scVI and scANVI, which use count data as input). On the Pancreas integration task, consisting of technical replicates obtained with different sequencing platforms, most methods performed well (**Fig. 3A**). In particular, Seurat CCA, STACAS and Seurat rPCA achieved high batch mixing (CiLISI) while preserving biological variance (cell type ASW). In this balanced scenario (**Fig. S1A**), cell type information does not appear to provide an advantage to semi-supervised methods: ssSTACAS obtained similar performance to unsupervised STACAS, scGen ranked below several unsupervised methods, and scANVI performed poorly and only marginally better than its unsupervised counterpart scVI.

**Figure 3:**
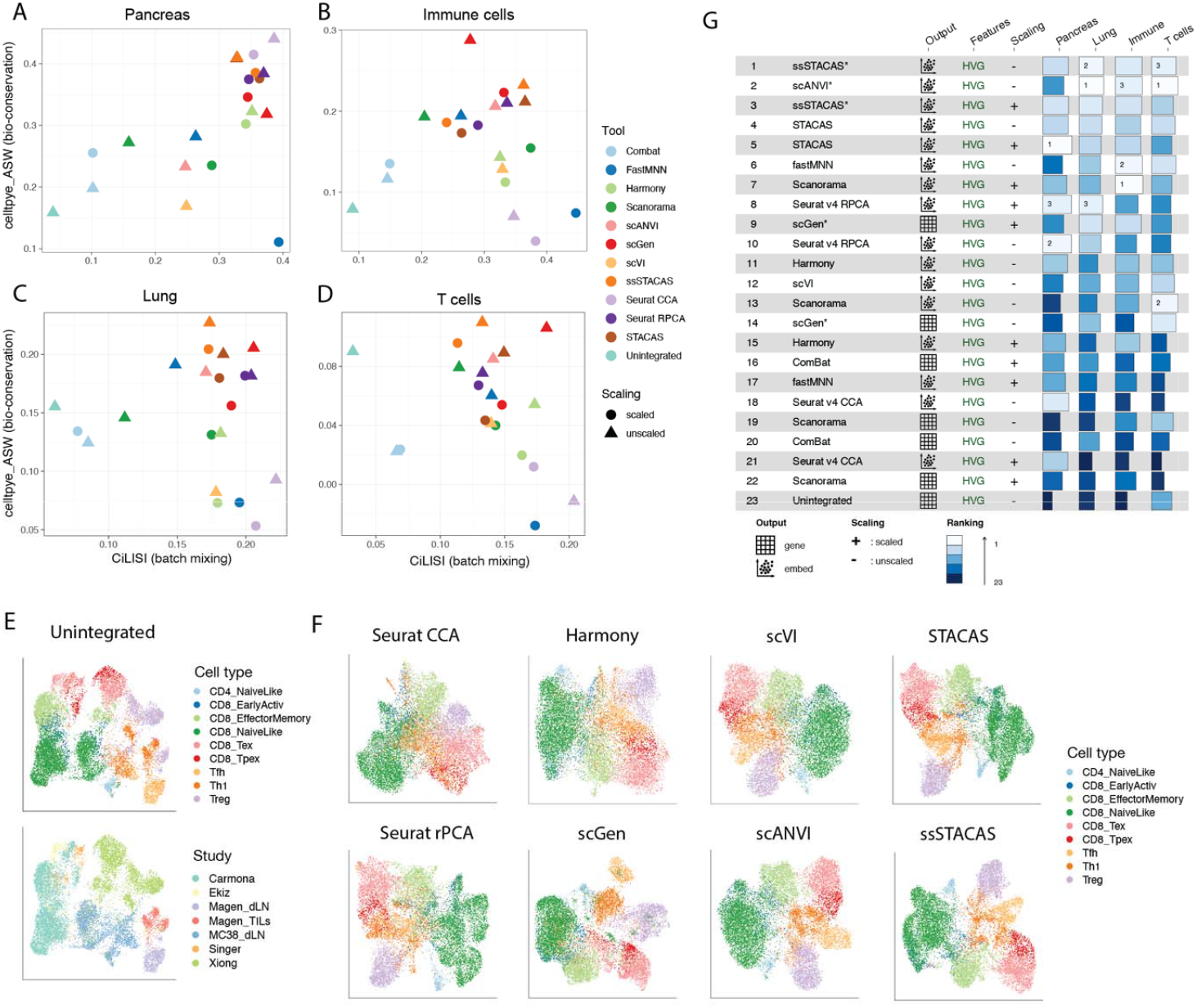
Integration performance for single-cell data integration tools over 4 different tasks. **A)** CiLISI (per cell type integration LISI, measuring cell type-aware batch mixing) vs. celltype_ASW (cell type average silhouette width, measuring preservation of biological variance) for several integration methods on the Pancreas integration task. **B)** CiLISI vs. celltype_ASW across methods on the Lung integration task. **C)** CiLISI vs. celltype_ASW across methods on the Immune cells integration task. **D)** CiLISI vs. celltype_ASW across methods on the T cells integration task. **E-F)** UMAP embeddings for the mouse T cell integration task, for unintegrated data colored by cell type (top) and by study of origin (bottom) **(E)** and for eight representative integration methods, colored by cell type **(F). G)** Global rankings of integration tools based on the weighted contribution of a broad panel of metrics both for preservation of biological variance (“bio-conservation”) and batch-correction, as proposed by Luecken *et al*. Supervised methods (ssSTACAS, scGen and scANVI) were provided noisy input labels (15% unknown and 20% shuffled labels). All experiments were performed with latent spaces of 50 dimensions. For alternative rankings using only CiLISI and celltype_ASW, or a different number of dimensions, see Fig. S2 and S3.

The Immune integration task comprises 10 datasets from 5 different studies corresponding to peripheral blood and bone marrow human samples, as compiled by Luecken et al.^12^. While most cell types are represented by multiple datasets, the composition is less balanced than in the Pancreas integration task (**Fig. S1B**) Here the semi-supervised methods scGen and ssSTACAS were the best tools in finding a trade-off between batch mixing and preservation of biological variance (**Fig. 3B**). Other methods such as Seurat CCA, scVI and Harmony, on the other hand, overcorrected batch effects and performed poorly in terms of cell type ASW (*cfr*. **Fig. 3B** and ‘overcorrected’ in **Fig. 2D**).

The Lung atlas represents a more challenging integration task, as it comprises samples from multiple human donors, covering different spatial locations^18^. Semi-supervised STACAS, and to a lesser degree its unsupervised version, successfully mitigated batch effects while preserving biological variance (**Fig. 3C**). Again, the widely used Seurat CCA, Harmony and scVI performed poorly in terms of cell type AWS, suggesting that these methods tend to overcorrect batch effects.

To evaluate the performance of STACAS in a setting with strong cell type imbalance and strong batch effects, we used the collection of T cell datasets from Andreatta et al.^19^. These contain seven datasets from six different studies covering tumor and lymph node samples, comprising studies with both CD4+ and CD8+ T cells (*MC38_dLN, Ekiz* and *Xiong*), only CD8+ T cells (*Carmona, Singer*) or only CD4+ T cells (*Magen_dLN* and *Magen_TILs*) (**Fig. S1D**). As with the Immune integration task, we observed that semi-supervised STACAS and scGen were the best performing tools, especially in terms of preservation of biological variance (**Fig. 3D-F**).

Globally, when considering cell type ASW and CiLISI across the four integration tasks, semi-supervised STACAS was the best performing method, followed by unsupervised STACAS and scGen (**Fig. S2**). When evaluating performance on a broad panel of metrics for preservation of biological variance (“bio-conservation”) and batch-correction, as in the original benchmark by Luecken et al., semi-supervised STACAS remains the best method, followed by the semi-supervised method scANVI (**Fig. 3G, S3**). These rankings remained consistent whether the latent spaces used 30 or 50 reduced dimensions (**Fig. S2-S3**). Across all integration tasks, using unscaled data was preferable to scaled data for the preservation of biological variance in terms of cell type silhouette coefficient (**Fig. S4**). In integration tasks with large cell type imbalance, methods that use prior cell type information were shown to better preserve biological variance compared to unsupervised methods. Based on these results, we recommend using prior cell type information to guide single-cell data integration tasks.

### Semi-supervised STACAS is robust to incomplete and noisy annotations

A potential risk of applying supervised or semi-supervised integration methods is overfitting the cell type labels provided as input. In real-life data integration scenarios, manual or automatic cell annotation may not allow providing cell type identities to all cells. Even when all cells can be annotated, these annotations might be wrong or inaccurate. Therefore, relying excessively on *a priori* cell type labels to force integration results may be undesirable. Instead, robust (semi) supervised integration methods should be tolerant to incomplete and incorrect input cell type information.

To evaluate the effect of incomplete or noisy cell type annotations on the performance of semi-supervised integration methods, we constructed alternative versions of the four collections of datasets used in our benchmark (Pancreas, Immune, Lung and T cell tasks) with increasing levels of shuffled or unknown cell type labels. We used these noisy labels as input for (semi) supervised integration, and the original labels to evaluate performance. As the percentage of shuffled labels increases from 0% to 100%, we observed an expected gradual drop in performance for a l supervised methods (**Fig. 4A**). However, scGen was considerably more sensitive to shuffled labels than ssSTACAS and scANVI, with a sharp drop in performance even with 10% or 20% of shuffled labels. Both ssSTACAS and scANVI were robust to noisy labels, but ssSTACAS obtained higher celltype ASW across all tasks. Similar results were observed when providing incomplete cell type labels as input to the three tools (0% to 100% ‘unknown’ labels) (**Fig. 4B**). These results suggest that scGen, when used as a batch-correction tool, is prone to overfitting on the provided input labels. In general, using the same cell type labels as input for supervised integration and to evaluate integration leads to over-optimistic performance assessment and should be avoided.

**Figure 4:**
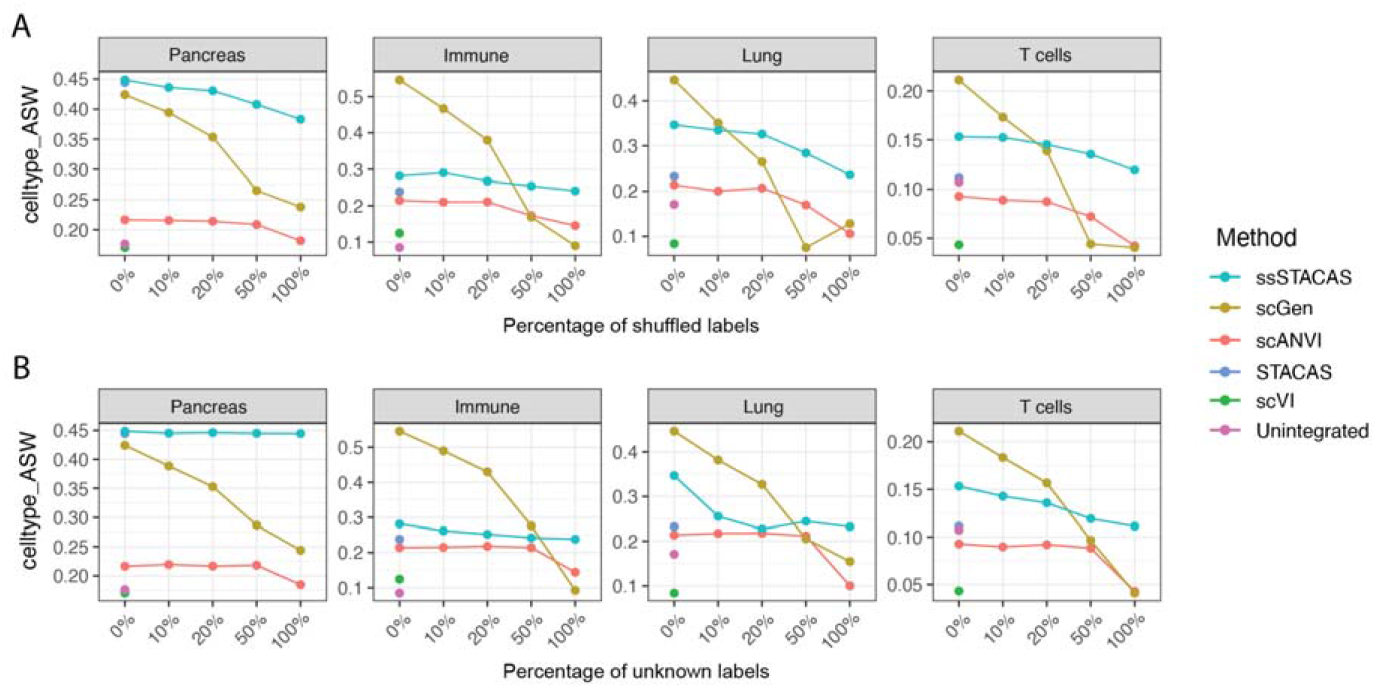
Effect of noisy or incomplete cell type annotations on data integration by supervised or semi-supervised methods. **A)** Cell type average silhouette coefficient (celltype_ASW) for 4 data integration tasks, using as input all cell type labels (original) or increasing levels of shuffled cell type labels (10% to 100%). **B)** Cell type average silhouette coefficient (celltype_ASW) for 4 data integration tasks, using as input all cell type labels (original) or increasing fractions of unknown cell type labels (10% to 100%). Unsupervised versions of ssSTACAS and scANVI (STACAS and scVI respectively) are included for reference.

In light of these results, we performed our main benchmark with 20% shuffled labels and 15% unknown labels. Because random shuffling of labels can, with some probability, swap identical labels, a 20% random shuffling results in practice in about 15% of actual shuffled labels of a different identity. This setup corresponds to an integration task where about 70% of cells are correctly annotated, 15% are wrongly annotated, and 15% are left unannotated. We believe this is a more reasonable setting for benchmarking than assuming the totality of true cell types can be known prior to integration. A modified version of the ‘scib’ pipeline that can account for incomplete or unknown annotations is available at https://github.com/carmonalab/scib-pipeline. Our results indicate that, while the use of (semi) supervised integration methods is highly recommended, it is critical to consider their robustness to missing and noisy input labels.

### Construction of a multi-study reference single-cell transcriptional map for human CD8 T cells

To date, most single-cell studies define cell states from scratch by dimensionality reduction, cell clustering and annotation. This approach is highly time-consuming and leads to inconsistent definitions across studies. Instead, the use of expert-curated reference maps to interpret single-cell data enables more consistent and faster cell state definitions ^19–21^. Building robust reference single-cell maps typically requires integrating multiple datasets from different studies and conditions.

To showcase the benefits of semi-supervised STACAS to generate a multi-study reference map, we applied it to integrate multiple human CD8+ T cell single-cell datasets. Starting from a publicly available collection of tumor-infiltrating lymphocyte scRNA-seq datasets (‘Utility’ collection), we identified 20 high-quality samples with a sufficiently large number of cells and high subtype diversity (see Methods). These samples amounted to 11,021 cells covering 7 different tumor types from multiple studies ^22–30^. Before dataset integration, we observed large batch effects between samples (**Fig. 6A**), as could also be quantified by a low CiLISI value (**Fig. 6B**). To leverage prior knowledge on CD8 T cell diversity, we defined scGate ^31^ gating models for six CD8 T cell subtypes based on well-established marker genes from literature (see Methods). This model allowed annotating individual datasets with partial labels and provided prior knowledge for semi-supervised integration (**Fig. 6A**).

**Figure 6:**
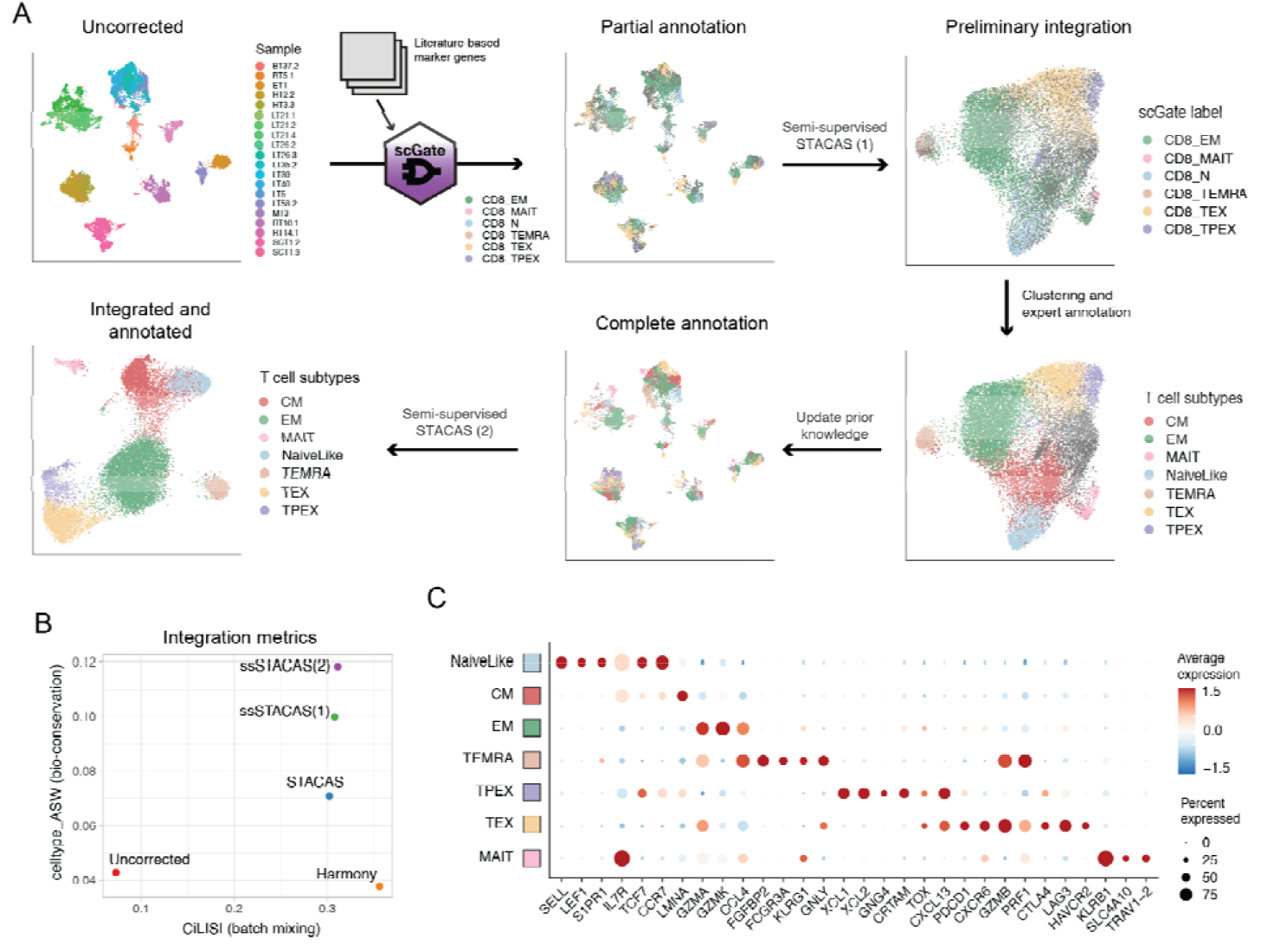
Construction of a multi-study reference map for human CD8 T cells with semi-supervised STACAS. **A)** Starting from 20 samples with large batch effects, partial cell type annotations were generated using the scGate package and literature-based marker genes. These labels were used as input to a first semi-supervised STACAS integration, allowing the mitigation of batch effect and the expert annotation of clusters corresponding to T cell subtypes. These new labels were used to update the prior knowledge of the original data, and used as input for a second semi-supervised STACAS integration, to define the final integrated space of CD8 T cell subtypes. **B)** Integration metrics for batch mixing (CiLISI) and biological variance preservation (celltype_ASW) for the uncorrected data, Harmony, unsupervised STACAS, semi-supervised STACAS on the initial partial annotation by scGate [ssSTACAS (1)] and semi-supervised STACAS on the updated annotations derived from the first integration [ssSTACAS (2)]. **C)** Average gene expression profiles (scaled by standard deviation) for a panel of marker genes on the seven T cell subtypes of the final integrated CD8 T cell map.

Upon semi-supervised STACAS integration, cells of the same type were clustered together, with simultaneous mitigation of batch effects (**Fig. 6A-B, Fig. S5A**). Compared to the uncorrected data, both batch mixing (CiLISI) and biological variance preservation (celltype ASW) were improved (**Fig. 6B**). Importantly, semi-supervised integration allowed higher celltype ASW compared to unsupervised STACAS integration, without a negative impact on batch mixing (**Fig. 6B**). Integration of these data by Harmony ^9^, one of the most widely used integration methods to date, resulted in high batch mixing but dramatic loss of biological variance (**Fig. 6B, Fig. S5A**), in agreement with the results from our benchmark. On the integrated space produced by this first integration (Semi-supervised STACAS (1)), we performed clustering and manually annotated the main T cell subtypes. We then went back to the original data with this updated, complete set of cell type annotations, and performed a new semi-supervised integration (Semi-supervised STACAS (2)) guided by the updated prior knowledge that was garnered from the first round of integration. The second integration allowed further improving celltype ASW while conserving good batch mixing (**Fig. 6A-B, Figure S5A**). This suggests a strategy to iteratively update and improve cell type annotations based first on prior knowledge, and secondly on the results of preliminary analyses.

The integrated single-cell map recapitulated with high resolution the known diversity of CD8 tumor-infiltrating T cells, as represented by seven different subtypes (**Fig. 6C** and **Fig. S5B**): Naive-like cells, characterized by the expression of transcription factors *TCF7* and *LEF1* and homing molecules *SELL, S1PR1* and *CCR7*; transcriptionally-related Central-memory (CM) cells, with high expression of *IL7R* and lower expression of other naive cells markers; Effector-memory (EM) cells, characterized by highest *GZMK* expression (which in the tumoral context have been referred to as ‘pre-dysfunctional’ ^32^); terminally differentiated effector cells (TEMRA), with high expression of *GNLY, PRF1*, multiple granzymes, *KLRG1* and *FCGR3A* (encoding CD16) ^33^; mucosal-associated invariant T cells (MAIT), characterized by the semi-invariant TCR chain *TRAV1-2* and expression of *KLRB1* ^34^; terminally exhausted effector (TEX) T cells, expressing cytotoxic molecules (*GZMB, PRF1*), multiple inhibitory receptors (e.g. *PDCD1, LAG3, CTLA4, HAVCR2*) and the exhaustion regulator *TOX* ^35,36^; and the elusive precursors of exhausted (TPEX) T cells, with co-expression of *TCF7, TOX* and *PDCD1*, and specific expression of chemokines *XCL1* and *XCL2* ^19,37,38^. These T cell subtypes showed variable frequency between samples (**Fig. S1E**), but displayed consistent expression profiles across studies, patients, and cancer types (**Fig. S5B**).

Finally, we evaluated whether STACAS can be used for large-scale integration of hundreds of samples and hundreds of thousands of cells. For large-scale integration (by default, more than 20 datasets), STACAS switches to a sequential integration strategy. Because sequential integration only requires calculation of integration anchors between one dataset at a time against a reference dataset, it is relatively undemanding in terms of computational resources.

All remaining datasets from the ‘Utility’ collection (265 datasets, totaling 553,077 cells) were sequentially integrated on the annotated reference of “seed” datasets (**Fig. 7A**). After recalculation of low-dimensional embeddings, we assigned cell type labels to all new cells by K-nearest neighbor classification using the reference, annotated cells. This resulted in an integrated collection of

**Figure 7:**
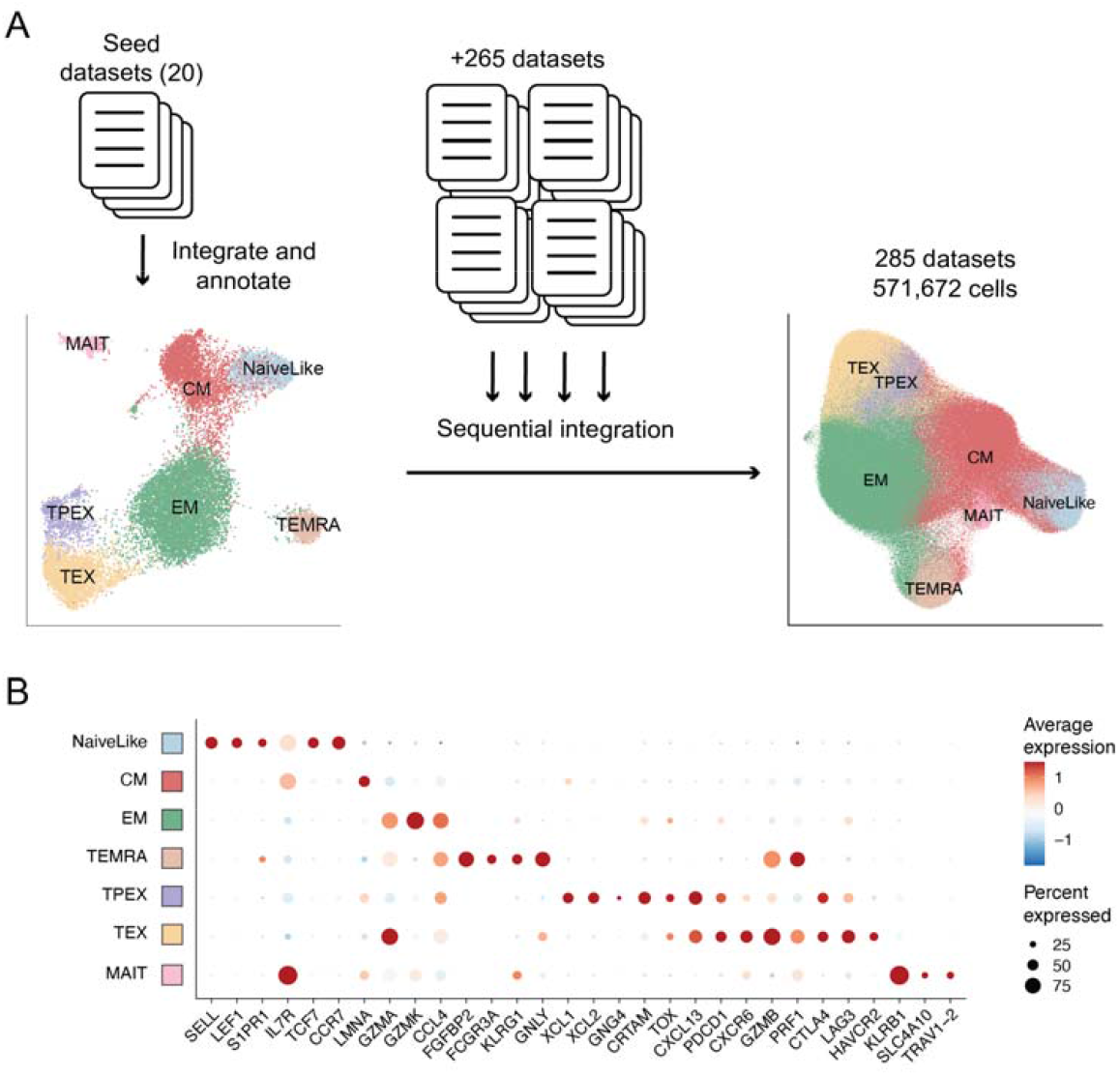
Large scale integration of hundreds of samples. **A)** A set of 20 high quality “seed” datasets was selected to perform an initial integration and annotation of cell types. All remaining datasets (265 datasets) were integrated using STACAS sequential integration mode over the “seed” integrated map. Cell labels for the new, unannotated cells were obtained by a kNN classifier that assigns the majority label from the top 20 nearest neighbors from the “seed” datasets. This results in an integrated set of > 500,000 cells from 285 samples. **B)** Average gene expression profile for a panel of marker genes, using all cells from 285 samples.

571,672 cells from 285 datasets, with average expression profiles for key T cell markers that matched expected profiles for the CD8 T cell subtypes (**Fig. 7B**). The integrated reference map, which can be directly applied to annotate additional datasets using ProjecTILs^19^, is available from figshare (https://doi.org/10.6084/m9.figshare.23608308) and can be explored interactively in SPICA^39^ at https://spica.unil.ch/refs/CD8T_human.

## Discussion

Integration of multiple single-cell transcriptomics datasets is a powerful approach to characterize cell diversity across tissues and conditions. Although decades of research have contributed extensive knowledge on cell markers that distinguish cell types, this information is typically not exploited for scRNA-seq data integration. In this work we propose a user-friendly tool that can take advantage of available prior information on expected cell types to obtain accurate data integration. In particular, we showed that even partial and imperfect annotations can be beneficial towards preserving biological variance while correcting for technical batch effects in integrated single-cell datasets.

In practice, some degree of prior knowledge on the cell type composition of biological samples is nearly always available. Marker genes from literature, as well as genes encoding markers commonly used in flow cytometry and immunohistochemistry are often available for many cell types. It is common practice in single-cell analyses to examine the expression of such markers to gain insights into cell cluster identity and provisionally identify cell types. We have previously shown that this task can be automated by computational tools such as scGate^31^. While complete and fine-grained annotation of cell types is challenging to achieve based only on a few marker genes, our results suggest that even incomplete prior knowledge is beneficial to data integration. Moreover, partial annotations can be complemented and refined by preliminary integration and annotation steps, suggesting a general strategy to iteratively update cell type labels and achieve improved dataset integration.

A recent benchmark^12^ showed that the two top performing methods for scRNA-seq integration were those that can make use of cell type labels information, scANVI^6^ and scGen^40^. However, the performance of these methods may have been overestimated by using the same cell type labels as input for supervised integration and for evaluation of integration performance. In particular, we observed that when introducing noisy or incomplete cell labels as input, the integration performance of scGen dropped significantly. In contrast, semi-supervised STACAS and scANVI were robust to noisy and incomplete annotations, maintaining high performance with levels of uncertainty commonly expected from single-cell datasets.

Quantifying batch effects in single-cell data is essential to assess integration quality and determine which integration methods and configurations work best in different scenarios. Commonly used batch-mixing metrics, such as LISI or Shannon entropy, are informative when evaluating batch effects between technical replicates, e.g. when there is no biological variability between samples. When integrating technical replicates, lower batch mixing between two technical replicates is a direct measurement of higher batch effects. However, most integration tasks with practical relevance involve distinct biological samples that display variation in cell type composition. In this case, a lower batch mixing does not necessarily imply higher batch effects. We argue that batch mixing metrics that neglect cell type information can overestimate batch effects between samples with large biological variance and underestimate batch effects in “overcorrected” data. This is particularly important when integrating datasets with significant cell type imbalance: in the absence of batch effects, cell type-agnostic batch-mixing metrics can increase when biological variation is removed, as cells of different type and batch are brought together as result of batch correction algorithms. In the context of benchmarks for integration methods, this translates into favoring single-cell integration methods that “overcorrect” and penalizing methods that preserve biological variance.

To overcome this issue we propose to use cell type-aware batch mixing metrics, such as CiLISI. Because CiLISI measures batch mixing of cells of the same type only, spurious removal of cell type variance is not associated with an artificial increase in batch mixing metrics. We note that Luecken et al.^12^ were also aware of this effect and previously suggested cell type-aware modifications to existing metrics (kBET and batch ASW). One obvious limitation of CiLISI and similar metrics is that it requires cell type annotations. As discussed above, i) the utility of cell type-agnostic batch mixing metrics is arguably very limited, and only relevant for integration of technical replicates, and ii) it is virtually always possible to provide some level of cell type annotation. Another limitation is that CiLISI cannot be calculated for cell types that are only present in one dataset. Our implementation excludes by default these cell types from calculation. We also note that, as LISI, CiLISI remains suboptimal when datasets have highly unequal numbers of cells (e.g. even in the absence of batch effects CiLISI would be lower than the optimal value of 1 that would be observed if all cell types were equally represented). Batch mixing metrics that are more robust to lopsided cell proportions are still lacking (see ref.^41^ for a systematic evaluation).

Our comprehensive benchmark using a reproducible pipeline showed that STACAS v2 consistently ranked as the top method across multiple integration tasks and using different combinations of integration metrics. The use of prior knowledge was particularly beneficial for the integration of datasets with large cell type imbalance, where semi-supervised methods more accurately preserved relevant biological variability. On a large collection of T cell samples from cancer patients, we demonstrated the feasibility of using STACAS to integrate hundreds of samples and over 500,000 cells while preserving the diversity of T cell subtypes contained within the samples. Combining semi-supervised integration with emerging approaches to summarize massive single-cell datasets, such as metacells^42,43^ and sketching^44^, together with on-disk, out-of-memory data representations (e.g. using DelayedArray objects^45^) will further increase scalability for organ atlas-level applications comprising millions of cells. Altogether, we propose STACAS as a first-line method for scRNA-seq data integration and encourage the broader use of prior cell type knowledge to guide integration and assess its quality.

## Methods

### STACAS integration method

STACAS is largely constructed on Seurat’s anchor-based integration approach. Given as input a list of normalized expression matrices, one for each dataset, the general aim is to determine batch-effect correction vectors between pairs of datasets; and by subtracting these correction vectors to calculate a corrected data matrix of “integrated” expression values. The corrected data matrix can be used for downstream analyses such as dimensionality reduction, unsupervised clustering and cell type annotation.

### Highly variable features and dimensionality reduction

The first step in STACAS integration is the calculation of highly variable genes. These are determined using the *FindVariableFeatures()* function from Seurat, but we also exclude certain classes of genes such as ribosomal, mitochondrial, and cell cycling genes that can have a large effect on the low-dimensional spaces without important contribution to cell type discrimination. These genes sets are available through the R package SignatuR (https://github.com/carmonalab/SignatuR; see e.g. https://carmonalab.github.io/STACAS.demo/STACAS.demo.html#notes-on-data-integration for an example). Additionally, genes with average log-normalized expression below (default 0.01) or above a threshold (default 3.0) are excluded from the highly variable genes. Consistently variable genes are then calculated as those found to be highly variable in multiple datasets, until reaching the desired number of genes (by default 1000). This feature selection step, also referred to as ‘sharedFeatures’ in the results, reduces the dimensionality of the data to a few hundred to few thousand genes, and ensures that batch effects are calculated on genes with informative variability. The dimensionality of the data is further reduced by Principal Components Analysis (PCA) from the set of consistently variable genes. Unlike the Seurat integration method, we do not rescale the data to zero mean and unit variance; we have previously shown how this step can mask important biological differences between datasets ^46^.

### Calculation and scoring of integration anchors

To determine integration anchors between pairs of datasets, we modified Seurat’s reciprocal PCA algorithm (rPCA) to find shared nearest neighbors and return the pairwise distance between anchors in rPCA space (rPCA_distance). These distances are used to weigh anchor contributions, in combination with Seurat’s shared nearest neighbor score (SNN_score) that quantifies the consistency of edges between cells in the same neighborhood of the SNN graph, using a geometric weighted sum:

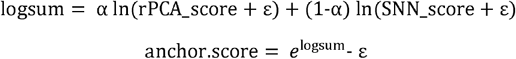

where ε is a small number (10^−6^) to avoid ln(0), and the rPCA distance is transformed to a score bound between 0 and 1 using:

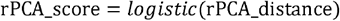

This procedure results in a set of integration anchors with associated weights, which can be used to estimate batch effects between pairs of datasets. The parameter α balances the contribution of the two scores, with α=0.8 by default. The score weighting scheme differs from previous versions of STACAS, where suboptimal anchors were directly filtered out based on the rPCA distance scores^46^. Re-weighting instead of filtering is more robust to parameter choices and avoids break cases where an insufficient number of integration anchors is retained after filtering, especially in the case of small datasets.

### Using prior knowledge to filter anchors

When available, prior information on cell types can be used to guide the integration by penalizing anchors composed of cells with inconsistent cell type labels. Cell type labels must be provided as a metadata column for each input object, and they can be incomplete, i.e. not all cells are required to have a label. Given a set of anchors calculated as described above and a set of cell type labels, the algorithm rejects (with probability = *label*.*confidence*) anchors composed of two cells with inconsistent labels. Cells without an annotation are never rejected at this step, generalizing to the unsupervised integration scenario when no labels are available. We recommend to only provide cell type labels for high-confidence associations to a cell type, and to leave the remaining cells as unlabeled (*NA* values). The outcome of this step is a subset of the previously calculated set of integration anchors, where anchors with inconsistent cell types have been removed.

### Integration guide trees

A crucial factor in the success of batch effect correction is the order of dataset integration. To this end, STACAS calculates a weight matrix that summarizes dataset-dataset similarity. For each pair of datasets, a similarity score is obtained by summing the combined anchor scores between the two datasets. On this similarity matrix, an integration tree is calculated by applying any of the clustering methods implemented in *hclust* from the *stats* package (by default ‘*ward*.*D2’*). Integration is initiated from the dataset with the highest sum of anchor scores against all other datasets; the rationale is that the dataset with the largest number of high scoring anchors should be the most “central”, with well-represented cell types, and a large number of cells. We note that, instead, Seurat’s integration trees are rooted by the largest dataset for each pair, regardless of the anchor scores.

### Performance metrics of data integration

The average silhouette width (ASW) quantifies the average distance of cells in a cluster compared to their distance to the closest of the other clusters. When applied to cell type labels (celltype ASW), it measures how well cells with the same cell type label are clustered together compared to other cell types in the dataset. We compute the celltype ASW with the ‘cluster’ package^47^ using euclidean distances in PC space, excluding cells with unknown cell type labels.

The Local Inverse Simpson Index (LISI) has been previously presented as a metric to quantify local batch mixing (iLISI) and cell type mixing (cLISI)^9^. Briefly, LISI metrics quantify the expected number of cells from different batches (or different subtypes) in a local neighborhood, with size determined by the perplexity parameter. We propose a modified version of iLISI that does not unfairly favor overcorrection by calculating it independently for each cell type. The new cell type-aware metric, called CiLISI, is then rescaled between 0 and 1 to make it comparable across integration tasks. In all experiments in this study we used a perplexity value of 30. Similarly, we redefined cLISI to vary between 0 and 1, where zero represents a random mix of cell types in all neighborhoods and one a perfect segregation of cell types; we call this quantity ‘normalized cLISI’. For normalized cLISI we set a perplexity value to twice the average number of cells per cell type per dataset.

We implemented these metrics and made them available as an R package at: https://github.com/carmonalab/scIntegrationMetrics.

### Comparison to other integration tools

Taking advantage of the previously published ‘scib’ pipeline for single cell integration benchmark pipeline^12^ we compared STACAS in unsupervised mode and semi-supervised mode (ssSTACAS) with 9 other integration tools: Combat^48^, Scanorama^4^, FastMNN^3^, Harmony^9^, Seurat v4 CCA and Seurat v4 rPCA^5^, scVI^49^, scANVI^6^ and scGen^40^. We conducted our benchmark on 4 integration tasks, as detailed next.

#### Integration tasks

The human pancreas atlas, the human immune cell atlas and the human lung atlas, collated by Luecken et al.^12^, were downloaded from figshare (https://doi.org/10.6084/m9.figshare.12420968.v8). We also included in our benchmark the mouse T cell atlas by Andreatta et al.^19^, available from figshare (https://doi.org/10.6084/m9.figshare.12478571). For these 4 collections of datasets, the preprocessed data (low quality cell filtered, raw and log-normalized counts as well as original cell type annotations) obtained from the previous studies were directly analysed with the ‘scib’ pipeline.

#### Integration procedure

We performed the 4 integration tasks as follows:

1. We used for all methods the same latent space dimensionality (D) for integration (e.g. number of principal components or dimensions of the reduced space or number of neurons in the bottleneck layer of autoencoders) and report here results for D=30 and D=50, the two most commonly used values in practice.
2. For Seurat-based methods (rPCA and CCA), we computed reduced dimensionalities for the integrated space directly in R, starting from the scaled corrected counts matrix and applying the *RunPCA()* Seurat function; the result is used as embedding output for the ‘scib’ pipeline.
3. All other methods were run with their default implementations as in the original benchmark; (ss)STACAS was run with the one-liner *RunSTACAS()* function and evaluated on the integrated PCA space it outputs.
4. For each integration task, we used the 2000 most variable genes across the different batches (automatically identified by the ‘scib’ pipeline) as integration features for all methods.
5. For each method, integration was performed both with or without a prior batch-aware scaling of the integration features (scaling + or – in the result summary), as implemented in the ‘scib’ pipeline.
6. Supervised integration with scANVI, scGEN and STACAS were conducted using noisy input cell type labels. First 15% of the original cell labels were set to ‘unknown’, then 20% of the remaining labels were shuffled randomly.

#### Integration metrics and ranking

We first compared the methods for their ability to remove batch effect while keeping distinct cell types separated using the CiLISI and celltype_ASW, respectively. We used the original cell type labels (i.e. the “true” labels without noise) to compute these metrics. Using the metric aggregation procedure by Luecken et al., we computed for each tool a combined score where CiLISI and cell type silhouette contribute with 60% and 40%, respectively. We calculated the mean score across all tasks to obtain a global score for each tool, which was used to compile a global ranking of the tools. Additionally, we also employed an alternative scoring scheme for tools, combining several metrics from the ‘scib’ pipeline: PCR batch, Batch ASW, graph iLISI, graph connectivity and kBET for the batch mixing score; NMI/cluster label, ARI cluster label, cell type silhouette, isolated label F1, isolated label silhouette, graph cLISI, cell cycle conservation and trajectory conservation (only for the immune cell atlas) for the bio-conservation score. We chose not to include HVG overlap as it can only be computed on the corrected matrix output, which is not available for all methods. As in the Luecken et al. benchmark, aggregation of metrics and ranking of tools was done by combining bio-conservation metrics and batch mixing metrics with a relative weight of 60% and 40%, respectively. Python methods producing a corrected matrix and a corrected integrated space were evaluated on both outputs, as in the original pipeline. R methods were only evaluated on their corrected integrated space (Harmony, Fastmnn) or computed in R after a scaling of their corrected feature matrix, as intended by the developers (Seurat, STACAS).

#### Robustness to noise of supervised methods

Supervised methods (scGen, ssSTACAS, scANVI) were further benchmarked using our pipeline by providing as input shuffled or unknown cell type annotation, with increasing levels of wrong/missing annotations (10, 20, 50 and 100 percent). In each scenario, the performance metrics (CiLISI and celltype_ASW) were evaluated on the “true” cell type labels.

#### UMAP of the integrated spaces

We calculated the two-dimensional UMAP for each method output (integrated space or corrected feature matrix) using the ‘scib’ pipeline. For corrected feature matrix outputs, a D-dimensional reduced space (D=30 or D=50) was calculated using a PCA of the corrected feature matrix, subsequently reduced to 2 dimensions using the UMAP approximation. For methods directly producing a D-dimensional integrated space, this was used directly as an input to calculate the UMAP representation.

### scGate prediction models

The scGate package ^31^ allows defining marker-based models for the annotation of cell types in single-cell datasets. For the annotation of human CD8 T cell subtypes, we used the collection of CD8_TIL models found at the scGate_models repository, (https://github.com/carmonalab/scGate_models, version v0.11). Briefly, the following literature-based signatures were used as gates to define subtypes: Naive-like cells: *LEF1+ CCR7+ TCF7+ SELL+ TOX-CXCL13-* ; Effector-memory: *GZMK+ CXCR3+; TEMRA: FCGR3A+ CX3CR1+ FGFBP2+*; Precursor-exhausted: *XCL1+ XCL2+ TOX+ GNG4+ CD200+*; Terminally exhausted: *TOX+ PDCD1+ LAG3+ TIGIT+ HAVCR2+*; MAIT: *TRAV1-2+ SLC4A10+*. Please refer to the models repository above for the complete combination of gates.

### Human CD8 TIL datasets

T cell scRNA-seq data were obtained from the ‘utility’ dataset collection (https://github.com/ncborcherding/utility), which collates data and harmonizes metadata from multiple studies and cancer types. Individual datasets were pre-processed using standard quality control, and homogenizing gene symbols according to Ensembl version 105. After filtering pure CD8 T cells, we applied UCell^50^ to remove cycling cells (UCell score > 0.1) and outliers in terms of interferon response (UCell score > 0.25). We applied the CD8_TIL scGate models (see section above) to obtain a preliminary annotation of subtypes in each dataset and estimate subtype diversity. Based on these annotations, we selected 20 “seed” datasets with a large number of cells and high subtype diversity, and used them for the construction of the reference map. For all versions of STACAS, we calculated 800 variable features, and further reduced the dimensionality of the data to 30 principal components. All other parameters were used as default values. The remaining samples in the ‘utility’ collection were then sequentially integrated using the default STACAS pipeline, specifying the “seed” integrated map as the base dataset using the *reference* parameter. Cell type labels were transferred from the annotated “seed” map to all remaining cells by k-nearest neighbor similarity using the *annotate*.*by*.*neighbors*() function implemented in STACAS. Reproducible code for these experiments can be found at: https://github.com/carmonalab/CD8_human_TIL_atlas_construction

### Generation of synthetic data

We generated five synthetic datasets with different levels of batch effect and batch size using the Splatter package, which applies a gamma-Poisson model to simulate gene expression distributions resembling real single-cell transcriptomics data^51^. Each dataset was composed of 1000 cells, and consisted of three batches and two cell types. The ‘batch0’ dataset had the three subtypes in equal proportions and zero batch effects (*batch*.*facLoc=0* in splatSimulate); ‘batchMild’ was generated by specifying *batch*.*facLoc=0*.*06* and *batch*.*facScale=0*.*06*; ‘batchStrong’ was generated by specifying *batch*.*facLoc=0*.*10* and *batch*.*facScale=0*.*10*; the ‘overcorrected’ dataset was simulated to have no differentially expressed genes between cell types by setting *de*.*facLoc=0* and *de*.*facScale=0*. In all datasets the two cell types were set to have equal proportions.

## Acknowledgements

We thank Nicholas Borcherding for compiling and maintaining the ‘Utility’ collection of tumor-infiltrating lymphocyte single-cell experiments with TCR sequencing data that were used in this study.

This work was supported by the Swiss National Science Foundation (SNF) Ambizione [180010] and by the ISREC foundation (to S.J.C).

## Ethics declarations

The authors declare no conflicts of interest.

## Code availability

STACAS is available as a R package at: https://github.com/carmonalab/STACAS

The snakemake pipeline that reproduces the results of our benchmark, based on the ‘scib’ pipeline by Luecken et al., is available at: https://github.com/carmonalab/scib-pipeline

The implementation of the performance metrics used in this work can be installed as a package from the following repository: https://github.com/carmonalab/scIntegrationMetrics

The code to construct, annotate and use the CD8 T cell reference map is available at: https://github.com/carmonalab/CD8_human_TIL_atlas_construction

## Supplementary Figures

**Figure S1:**
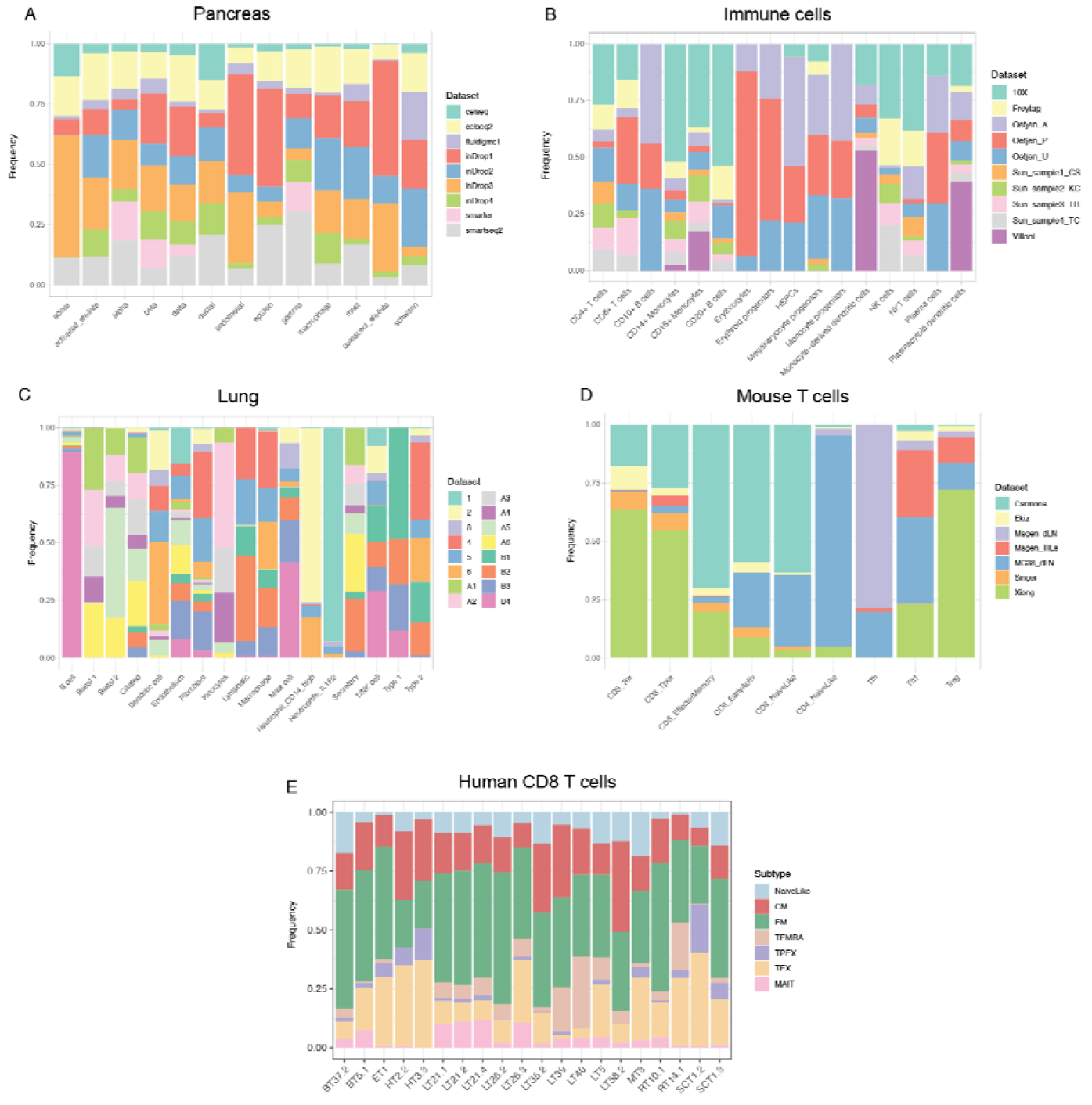
Dataset composition for individual cell types in 4 integration tasks. **A)** The Pancreas dataset is “balanced” in terms of cell type composition, with most cell types represented in the majority of datasets. **B)** In the Immune cell dataset several cell types are only represented in 3 of the 10 datasets. **C)** The lung integration task is challenging because it comprises samples from multiple human donors, covering different spatial locations, resulting in an imbalanced composition. **D)** The T cell dataset has also high cell type imbalance, with several cell type being represented by only one or few datasets. **E)** Cell type composition of the 20 samples used to construct the human CD8T map.

**Figure S2:**
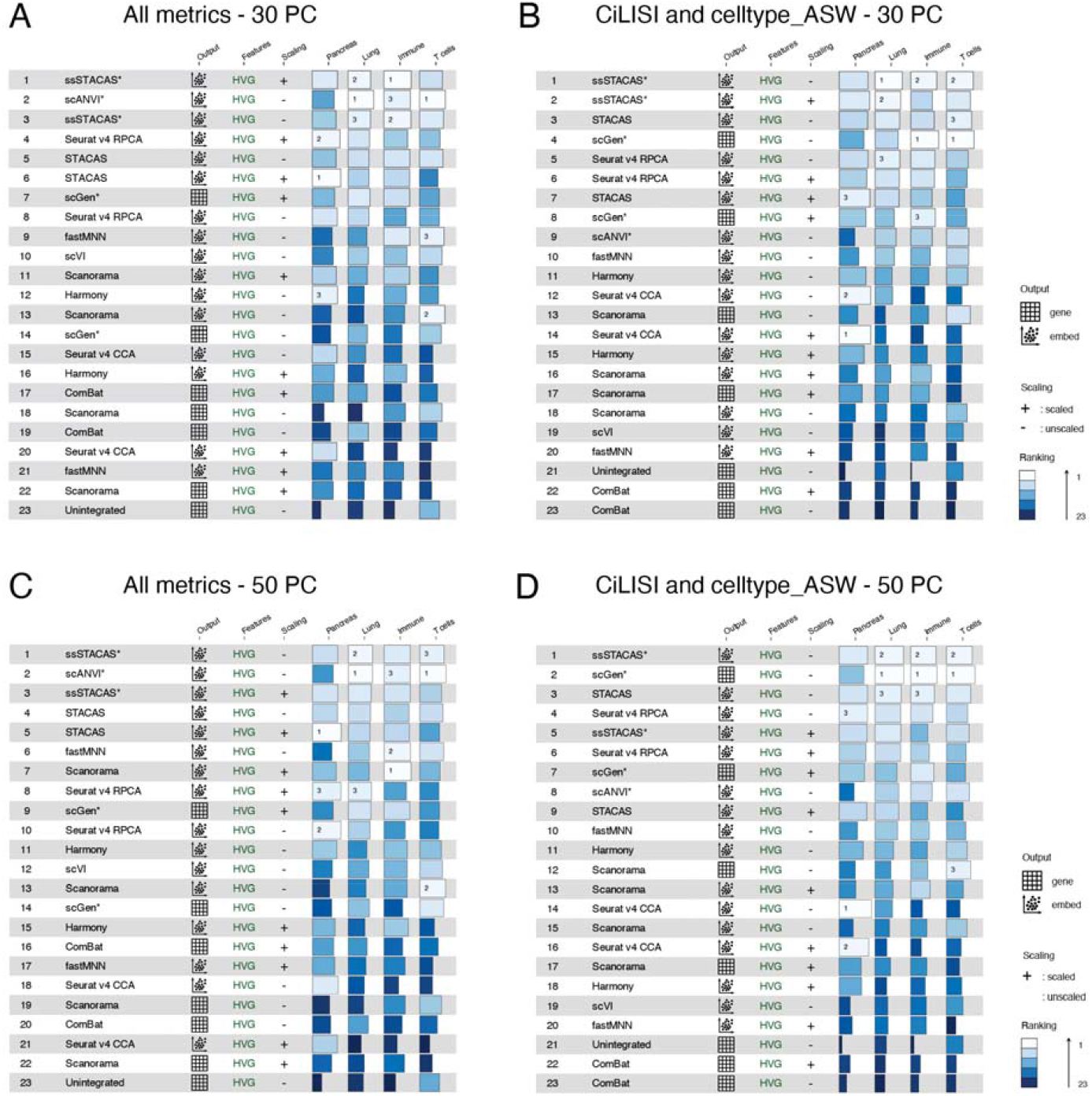
Global rankings for method performance using different metrics and dimensionality reduction sizes. **A)** Global rankings of integration tools based on the weighted contribution of multiple metrics proposed by Luecken et al., using 30-dimensional latent space (e.g. number of principal components for dimensionality reduction). **B)** Global rankings of integration tools based on the weighted contribution of CiLISI (quantifying batch mixing) and celltype_ASW (quantifying preservation of biological variance), and 30-dimensions latent space. **C)** As A) but using 50-dimensions latent space. **D)** As B) but using 50-dimensions latent space.

**Figure S3:**
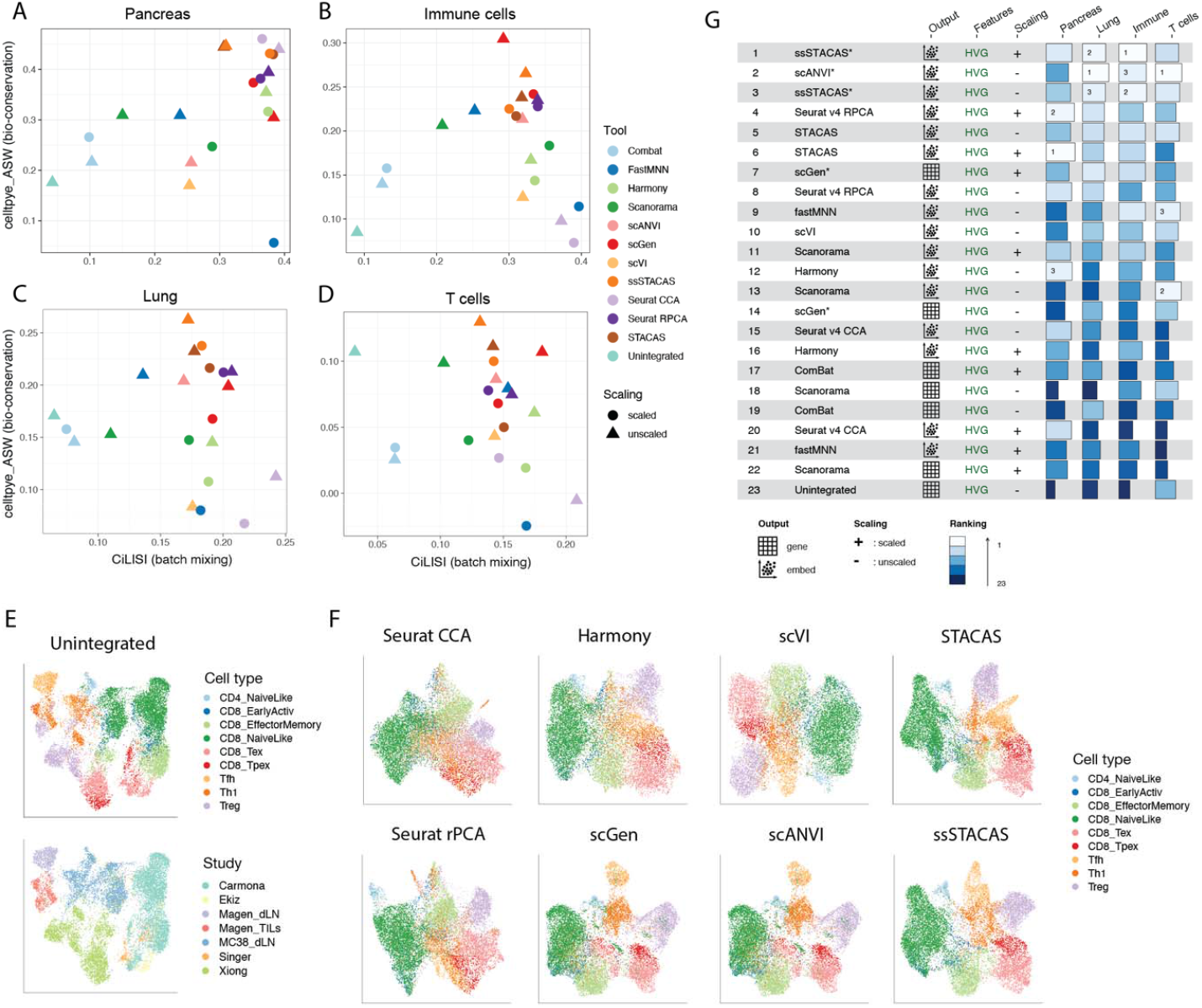
Integration performance for single-cell data integration tools over 4 different tasks, using a 30-dimensional latent space. **A)** CiLISI (per cell type integration LISI, measuring cell type-aware batch mixing) vs. celltype_ASW (cell type average silhouette width, measuring preservation of biological variance) for several integration methods on the Pancreas integration task. **B)** CiLISI vs. celltype_ASW across methods on the Lung integration task. **C)** CiLISI vs. celltype_ASW across methods on the Immune cells integration task. **D)** CiLISI vs. celltype_ASW across methods on the T cells integration task. **E-F)** UMAP embeddings for the mouse T cell integration task, for unintegrated data colored by cell type (top) and by study of origin (bottom) **(E)** and for eight representative integration methods, colored by cell type **(F). G)** Global rankings of integration tools based on the weighted contribution of a broad panel of metrics both for preservation of biological variance (“bio-conservation”) and batch-correction, as proposed by Luecken *et al*.

**Figure S4:**
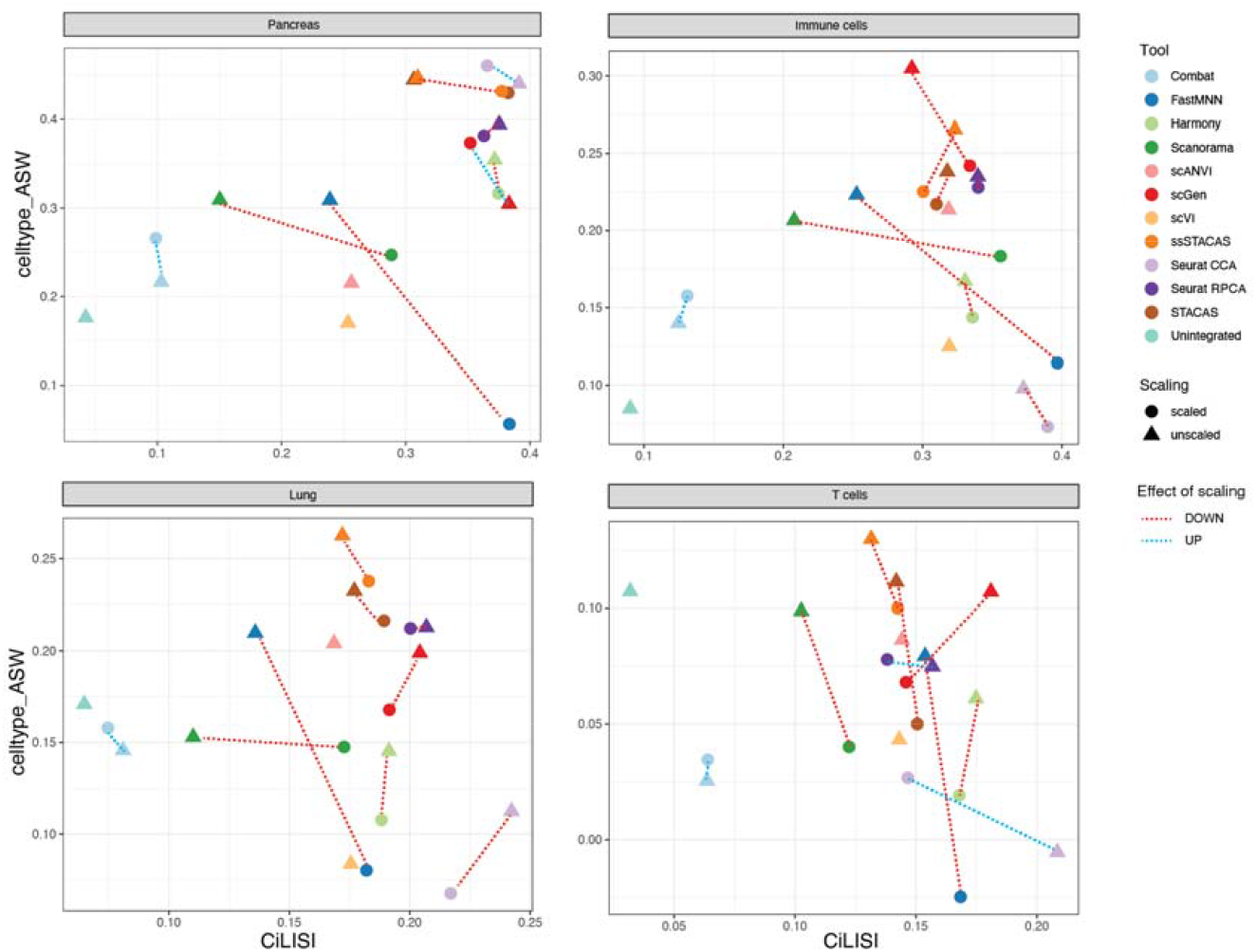
Effect of scaling on integration performance. CiLISI (batch mixing) versus celltype_ASW (preservation of biological variance) for indicated computational tools over 4 integration tasks. Dashed lines connect performance coordinates for the same tool with or without prior data scaling; blue lines indicate tools for which scaling increased celltype_ASW, red lines indicate tools for which scaling had negative effect on celltype_ASW.

**Figure S5:**
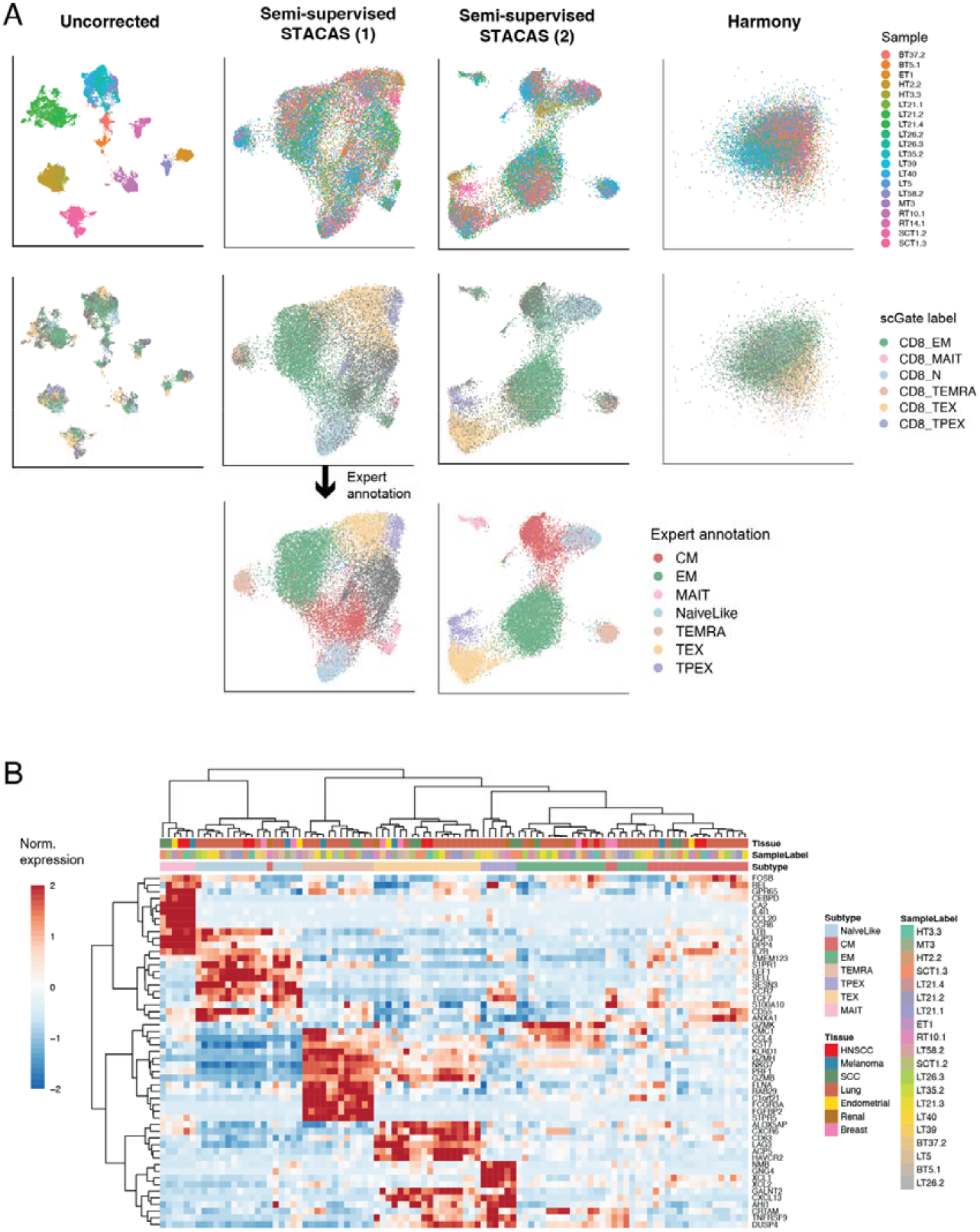
Construction of a reference map for CD8 T cells using semi-supervised STACAS. **A)** UMAP embeddings for unintegrated data, semi-supervised STACAS guided by scGate predicted cell types (1), semi-supervised STACAS guided by updated labels after expert annotation (2), and the Harmony integration tool. Cells are colored by sample of origin (first row), scGate predicted annotation (second row) and by expert annotation (third row). **B)** Average normalized RNA expression for differentially expressed genes in individual subtype-sample combinations of the CD8 T reference map. For all subtypes with at least 30 cells in a sample, the average normalized RNA expression was centered and rescaled by the standard deviation by gene. Profiles are shown for genes that were differentially expressed (log-fold change > 0.5) consistently in least 80% of the datasets. Hierarchical clustering using the Ward D2 algorithm shows that expression profiles are grouped largely by T cell subtype rather than by study or tissue of origin.

## References

1. Argelaguet, R., Cuomo, A. S. E., Stegle, O. & Marioni, J. C. Computational principles and challenges in single-cell data integration. Nat Biotechnol 39, 1202–1215 (2021).

2. Tran, H. T. N. et al. A benchmark of batch-effect correction methods for single-cell RNA sequencing data. Genome Biology 21, 12 (2020).

3. Haghverdi, L., Lun, A. T. L., Morgan, M. D. & Marioni, J. C. Batch effects in single-cell RNA-sequencing data are corrected by matching mutual nearest neighbors. Nat Biotechnol 36, 421–427 (2018).

4. Hie, B., Bryson, B. & Berger, B. Efficient integration of heterogeneous single-cell transcriptomes using Scanorama. Nature Biotechnology 37, 685–691 (2019).

5. Stuart, T. et al. Comprehensive integration of single-cell data. Cell 177, 1888–1902.e21 (2019).

6. Xu, C. et al. Probabilistic harmonization and annotation of single-cell transcriptomics data with deep generative models. Mol Syst Biol 17, e9620 (2021).

7. Azizi, E. et al. Single-Cell Map of Diverse Immune Phenotypes in the Breast Tumor Microenvironment. Cell 174, 1293–1308.e36 (2018).

8. Büttner, M., Miao, Z., Wolf, F. A., Teichmann, S. A. & Theis, F. J. A test metric for assessing single-cell RNA-seq batch correction. Nat Methods 16, 43–49 (2019).

9. Korsunsky, I. et al. Fast, sensitive and accurate integration of single-cell data with Harmony. Nature Methods 16, 1289–1296 (2019).

10. Rand, W. M. Objective criteria for the evaluation of clustering methods. Journal of the American Statistical association 66, 846–850 (1971).

11. Rousseeuw, P. J. Silhouettes: a graphical aid to the interpretation and validation of cluster analysis. Journal of computational and applied mathematics 20, 53–65 (1987).

12. Luecken, M. D. et al. Benchmarking atlas-level data integration in single-cell genomics. Nat Methods 19, 41–50 (2022).

13. Cao, Y. et al. scDC: single cell differential composition analysis. BMC Bioinformatics 20, 721 (2019).

14. Lun, A. T. L., Richard, A. C. & Marioni, J. C. Testing for differential abundance in mass cytometry data. Nat Methods 14, 707–709 (2017).

15. Maan, H. et al. The differential impacts of dataset imbalance in single-cell data integration. 2022.10.06.511156 Preprint at https://doi.org/10.1101/2022.10.06.511156 (2022).

16. Richards, L. M. et al. A comparison of data integration methods for single-cell RNA sequencing of cancer samples. 2021.08.04.453579 https://www.biorxiv.org/content/10.1101/2021.08.04.453579v1 (2021) doi:10.1101/2021.08.04.453579.

17. Sikkema, L. et al. An integrated cell atlas of the lung in health and disease. Nature Medicine 1–15 (2023).

18. Vieira Braga, F. A. et al. A cellular census of human lungs identifies novel cell states in health and in asthma. Nat Med 25, 1153–1163 (2019).

19. Andreatta, M. et al. Interpretation of T cell states from single-cell transcriptomics data using reference atlases. Nature Communications 12, 2965 (2021).

20. Kang, J. B. et al. Efficient and precise single-cell reference atlas mapping with Symphony. Nat Commun 12, 5890 (2021).

21. Lotfollahi, M. et al. Mapping single-cell data to reference atlases by transfer learning. Nat Biotechnol 1–10 (2021) doi:10.1038/s41587-021-01001-7.

22. Bassez, A. et al. A single-cell map of intratumoral changes during anti-PD1 treatment of patients with breast cancer. Nature Medicine 27, 820–832 (2021).

23. Wu, T. D. et al. Peripheral T cell expansion predicts tumour infiltration and clinical response. Nature 579, 274–278 (2020).

24. Eberhardt, C. S. et al. Functional HPV-specific PD-1+ stem-like CD8 T cells in head and neck cancer. Nature 597, 279–284 (2021).

25. Caushi, J. X. et al. Transcriptional programs of neoantigen-specific TIL in anti-PD-1-treated lung cancers. Nature 1–7 (2021) doi:10.1038/s41586-021-03752-4.

26. Liu, B. et al. Temporal single-cell tracing reveals clonal revival and expansion of precursor exhausted T cells during anti-PD-1 therapy in lung cancer. Nat Cancer 3, 108–121 (2022).

27. Banta, K. L. et al. Mechanistic convergence of the TIGIT and PD-1 inhibitory pathways necessitates co-blockade to optimize anti-tumor CD8+ T cell responses. Immunity 55, 512–526.e9 (2022).

28. Pauken, K. E. et al. Single-cell analyses identify circulating anti-tumor CD8 T cells and markers for their enrichment. J Exp Med 218, e20200920 (2021).

29. Krishna, C. et al. Single-cell sequencing links multiregional immune landscapes and tissue-resident T cells in ccRCC to tumor topology and therapy efficacy. Cancer Cell 39, 662–677.e6 (2021).

30. Yost, K. E. et al. Clonal replacement of tumor-specific T cells following PD-1 blockade. Nature Medicine 25, 1251–1259 (2019).

31. Andreatta, M., Berenstein, A. J. & Carmona, S. J. scGate: marker-based purification of cell types from heterogeneous single-cell RNA-seq datasets. Bioinformatics btac141 (2022) doi:10.1093/bioinformatics/btac141.

32. van der Leun, A. M., Thommen, D. S. & Schumacher, T. N. CD8+ T cell states in human cancer: insights from single-cell analysis. Nat Rev Cancer 20, 218–232 (2020).

33. Martos, S. N. et al. Single-cell analyses identify dysfunctional CD16+ CD8 T cells in smokers. Cell Rep Med 1, 100054 (2020).

34. Godfrey, D. I., Koay, H.-F., McCluskey, J. & Gherardin, N. A. The biology and functional importance of MAIT cells. Nature immunology 20, 1110–1128 (2019).

35. Blank, C. U. et al. Defining ‘T cell exhaustion’. Nature reviews. Immunology 1–10 (2019) doi:10.1038/s41577-019-0221-9.

36. Jin, H.-T. et al. Cooperation of Tim-3 and PD-1 in CD8 T-cell exhaustion during chronic viral infection. Proceedings of the National Academy of Sciences 107, 14733–14738 (2010).

37. Held, W., Siddiqui, I., Schaeuble, K. & Speiser, D. E. Intratumoral CD8+ T cells with stem cell–like properties: Implications for cancer immunotherapy. Science Translational Medicine 11, eaay6863 (2019).

38. Kallies, A., Zehn, D. & Utzschneider, D. T. Precursor exhausted T cells: key to successful immunotherapy? Nature Reviews Immunology 20, 128–136 (2020).

39. Andreatta, M., David, F. P. A., Iseli, C., Guex, N. & Carmona, S. J. SPICA: Swiss portal for immune cell analysis. Nucleic Acids Res 50, D1109–D1114 (2022).

40. Lotfollahi, M., Wolf, F. A. & Theis, F. J. scGen predicts single-cell perturbation responses. Nat Methods 16, 715–721 (2019).

41. Lütge, A. et al. CellMixS: quantifying and visualizing batch effects in single-cell RNA-seq data. Life Sci Alliance 4, e202001004 (2021).

42. Baran, Y. et al. MetaCell: analysis of single-cell RNA-seq data using K-nn graph partitions. Genome Biology 20, 206 (2019).

43. Bilous, M. et al. Metacells untangle large and complex single-cell transcriptome networks. BMC Bioinformatics 23, 336 (2022).

44. Hie, B., Cho, H., DeMeo, B., Bryson, B. & Berger, B. Geometric Sketching Compactly Summarizes the Single-Cell Transcriptomic Landscape. Cell Systems 8, 483–493.e7 (2019).

45. Pagès, H. HDF5Array: HDF5 backend for DelayedArray objects. R package version (2020).

46. Andreatta, M. & Carmona, S. J. STACAS: Sub-Type Anchor Correction for Alignment in Seurat to integrate single-cell RNA-seq data. Bioinformatics 37, 882–884 (2021).

47. Maechler, M., Rousseeuw, P., Struyf, A., Hubert, M. & Hornik, K. Cluster: cluster analysis basics and extensions. (2012).

48. Zhang, Y., Parmigiani, G. & Johnson, W. E. ComBat-seq: batch effect adjustment for RNA-seq count data. NAR genomics and bioinformatics 2, lqaa078 (2020).

49. Lopez, R., Regier, J., Cole, M. B., Jordan, M. I. & Yosef, N. Deep generative modeling for single-cell transcriptomics. Nat Methods 15, 1053–1058 (2018).

50. Andreatta, M. & Carmona, S. J. UCell: Robust and scalable single-cell gene signature scoring. Computational and Structural Biotechnology Journal 19, 3796–3798 (2021).

51. Zappia, L., Phipson, B. & Oshlack, A. Splatter: simulation of single-cell RNA sequencing data. Genome Biol 18, 174 (2017).

